# Aerosol delivery–based serial femtosecond crystallography

**DOI:** 10.64898/2026.04.21.719866

**Authors:** Yoonhee Kim, Faisal H. M. Koua, Raphael de Wijn, Lena Worbs, Robin Schubert, Sravya Kantamneni, E Juncheng, Egor Sobolev, Diogo Melo, Oleksii Turkot, Agnieszka Wrona, Marco Kloos, Chenxi Wei, Qing Xie, Adam Round, Jayanath Koliyadu, Marcin Sikorski, Romain Letrun, Katerina Doerner, Huijong Han, Joachim Schulz, Adrian P. Mancuso, Tokushi Sato, Richard Bean, Johan Bielecki, Chan Kim

## Abstract

Serial femtosecond crystallography (SFX) has revolutionised structural biology by enabling direct visualization of conformational dynamics of biomacromolecules at near-physiological temperatures. However, conventional SFX sample delivery at X-ray free-electron laser (XFEL) facilities introduces a significant X-ray scattering background, which may limit the data quality and resolution. Here, we introduce an aerosol delivery-based method that drastically reduces the background scattering by minimising the liquid environment surrounding the nanocrystals. We validate our method by solving the structure of *Cydia pomonella* granulovirus nanocrystals at 1.9 Å resolution, achieving orders-of-magnitude lower background scattering compared to a liquid jet-based method. Structural comparison with the liquid jet-based model revealed similar overall structure, suggesting that the native structure is largely preserved despite dehydration during aerosolisation. Our method enables efficient SFX studies, particularly pump-probe time-resolved SFX on protein nanocrystals with enhanced signal-to-noise ratios, as well as high-throughput small molecule SFX (smSFX) applications for pharmaceuticals and functional materials.

---

Serial femtosecond crystallography (SFX) [1–5] is one of the most active research methods in X-ray free-electron laser (XFEL) facilities [6–10]. It enables tracking structural and conformational changes of biomacromolecules in crystals at near-physiological temperature and pressure, with high spatiotemporal resolution, while minimising observable radiation damage [11]. Delivery of protein crystals for SFX experiments can be carried out by various liquid injection systems [12], e.g. gas-dynamic virtual nozzles (GDVNs) [13], double flow-focusing nozzles (DFFNs) [14], a high viscosity extrusion (HVE) injector [15, 16], and drop-on-demand (DoD) [17–20] systems. Sample delivery systems at high-repetition-rate XFELs, with an intra-train repetition rate of kHz to MHz regime, where each intense X-ray pulse destroys the sample, require a continuous and rapid supply of fresh samples [21, 22]. Currently used fixed-target and HVE methods are not well suited for the high-repetition-rate facilities.

High X-ray scattering background signals from liquid or lipidic cubic phase injection system, however, is one major limiting factor in accurate measurements of intensities to explore the dynamics of macromolecules in nanocrystals [23]. Fixed target sample delivery systems, such as silicon nitride or polymer membranes [24–27], can provide lower background noise; nonetheless, they are applicable only to low-repetition-rate XFELs, operating at less than kHz. To minimise the background scattering signal, the aerosol injection system [28–30] was developed and used primarily for single-particle X-ray diffraction imaging (SPI) experiments [31–35], where samples are delivered as aerosolised droplets with reduced water-layer.

A well-optimised aerosolised particle beam injection system [28–30] has advantages at XFEL facilities, especially at high-repetition-rate XFELs, in exploring weakly scattering biomacromolecules and nanocrystals. An earlier SFX experiment on aerosolised *Cydia pomonella* granulovirus (CpGV) reported by Awel *et al.* [36] demonstrated that the background scattering signal can be reduced by three orders of magnitude in comparison with typical liquid injection systems, though measured only a limited number of diffraction patterns, resulting in a partial dataset. In their experiment, isolated CpGV nanocrystals were delivered using a nebulisation chamber, transport tube, and convergent nozzle [37]. Here, we employ an aerodynamic lens stack (ALS) to generate and focus the nanocrystal beam, demonstrating the first successful SFX study of aerosolised single CpGV granulin nanocrystals with orders of magnitude reduced background.

## Results

### Aerosol SFX setup

We developed an aerosol SFX experimental setup illustrated in Fig. 1. The system consists of a nebulization chamber where the aerosol is generated, a differential pumping section to reduce the gas pressure, and an ALS that produces a focused beam of nanocrystals in the interaction region. The aerosol sample delivery system used here has previously been proven to be suitable for SPI experiments on individual nanoparticles [38]. In our experimental setup, aerosol particles are generated from a GDVN or a DFFN, operating at liquid flow rates of 2 − 5 *µ*l · min^−1^ (see Supplementary Table 1 for details).

**Fig. 1:**
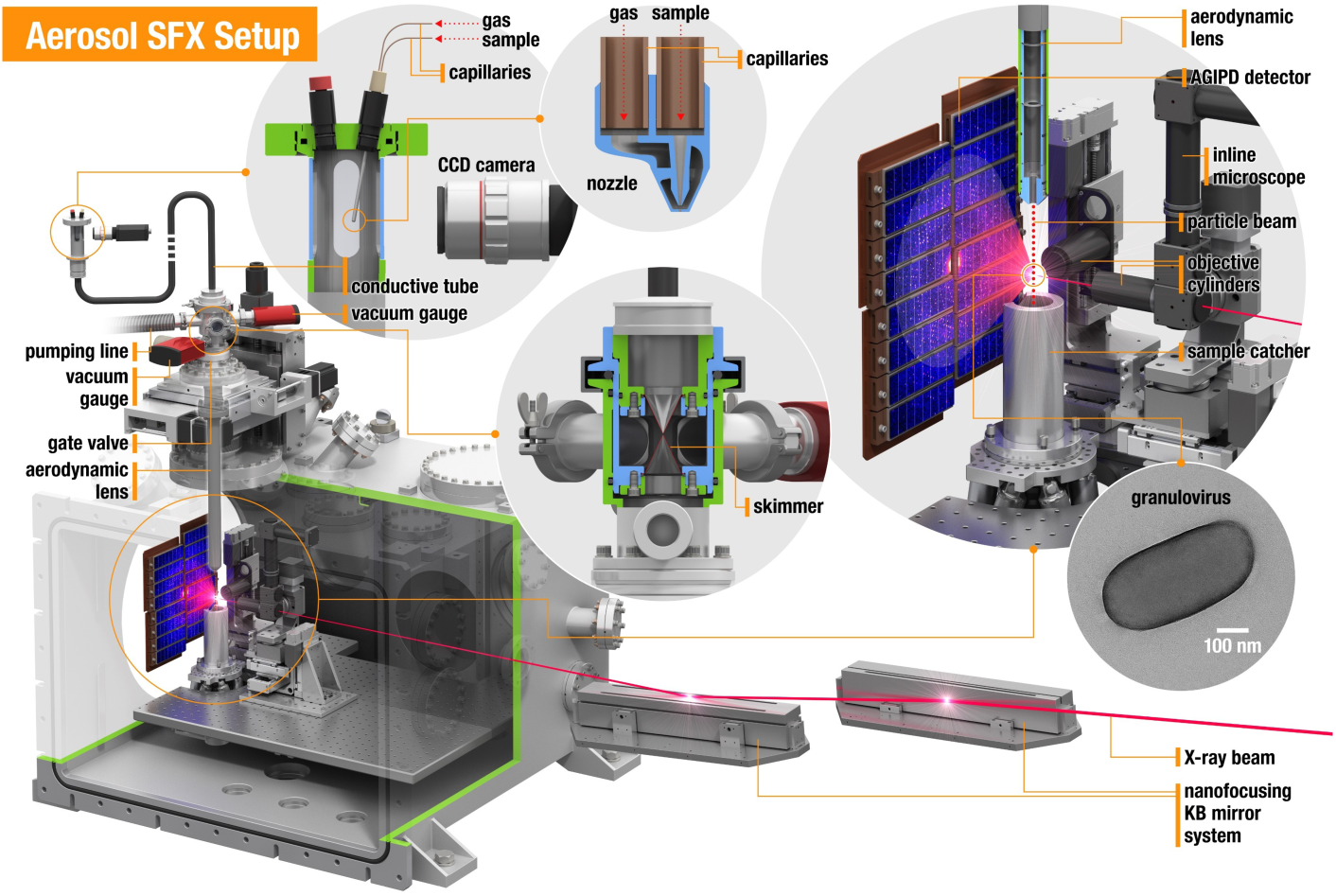
Aerosol SFX setup at the SPB/SFX instrument. Sample droplets were generated using a GDVN or DFFN and transported to the sample X-ray interaction point by passing through a transport tube, a skimmer, and an ALS. Excess gas was removed at the skimmer before the sample entered the main chamber, which was maintained under high vacuum during the experiment. The used samples were collected by a catcher located below the interaction region in order to prevent contamination inside the chamber. The X-ray beam was focused by nano-focusing KB mirrors.

Assuming a 2 *µ*m droplet diameter at the highest flow rate limit, the CpGV occupancy in the droplets follows a Poisson distribution (*λ*=1.05), resulting in 35 % of droplets being empty, 37 % containing single nanocrystals, 19 % containing two, and the rest of the droplets containing more than two. The droplet generation rate is in the range of 2 × 10^7^ s^−1^.

In our setup, helium gas and the generated droplets containing CpGV nanocrystals, pass through a ∼ 1.5 m long conductive tube into the differential pumping section in a rough vacuum environment—most likely driving the dehydration of the sample. Dehydration of the droplets can result in clusters of CpGV nanocrystals and may lead to an increased rate of multi-crystals per droplet. The nanocrystal droplets arrive in the differential pumping section consisting of a nozzle and skimmer, where most of the helium is removed, before entering the ALS. While passing through the ALS, based on our simulation, particles are focused at the interaction region and accelerated to nearly identical velocities (Supplementary Fig. 1 a,b).

The nanocrystal beam is focused, with a calculated minimum width (less than 10 *µ*m) at a distance of 4.2 mm from the exit of the ALS. The calculated speed of the nanocrystals is at least 43 m · s^−1^ at 1.2 mbar and 58 m · s^−1^ at 1.9 mbar, which is fast enough to replace an X-ray pulse-exposed nanocrystal before the next pulse at a repetition rate of 1.128 MHz, commonly used at European XFEL (Supplementary Fig. 1 b,c). The speed of the nanocrystals depends on the mass and the pressure before the ALS and can be increased by lowering the mass of the crystal or increasing the pressure, allowing for even higher X-ray repetition rates without increasing sample consumption. As shown in Fig. 2, the aerosol delivery method demonstrates both a lower sample consumption rate and the unique capability to operate at 4.5 MHz—the highest current intra-train repetition-rate—surpassing other delivery methods and making it a potential method of choice for future high repetition rate facilities.

**Fig. 2:**
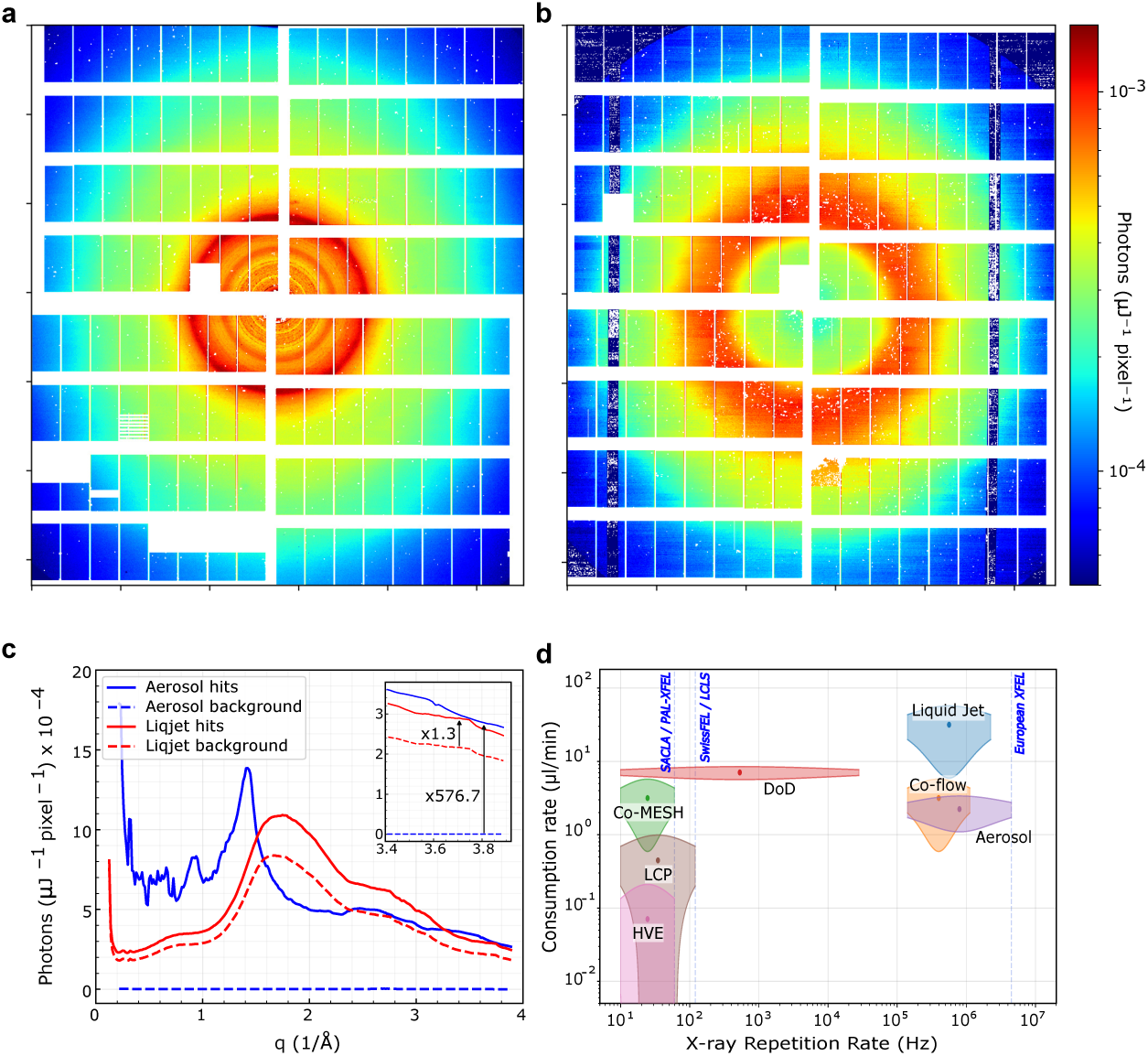
Background scattering signal and consumption rate of aerosol delivery method. Bragg peaks of CpGVs nanocrystals delivered using (a) aerosol delivery method and (b) liquid jet. (c) The azimuthally averaged radial distributions of aerosol and liquid jet delivery methods plotted as a function of resolution after normalization using the pulse energy. The SNR at ≈ 3.8 Å^−1^ is shown in the inlet. (d) The consumption rates (assuming an average concentration of ∼1× 10^10^ crystals per ml) of the aerosol delivery in comparison with different delivery methods as function of their X-ray repetition rates capability at the different facilities. The plot represents the operating rate for each method as defined by the minimum and maximum consumption rates and the lowest and maximum reachable X-ray repetition rates.

To validate the aerosol SFX setup, we performed two independent experiments at 6 keV and another one at 9.3 keV. In these experiments, we achieved hit rates of ∼0.03 %, five times higher than the convergent-nozzle aerosol injector setup [36], which is slightly less than, but still comparable with, that of typical SPI experiments with biological samples [34, 35, 39]. The aerosol sample delivery provides a quasi-background-free method for delivering nanocrystals into the interaction region compared with liquid-jet based SFX (Fig. 2a,b). Figure 2c shows azimuthally averaged diffraction and background scattering signals (up to q-range = 3.85 Å^−1^), for aerosol and liquid-jet sample delivery systems, which were integrated from 4200 diffraction patterns pre-normalized to the incident pulse energy. The results demonstrate that aerosol SFX exhibits a substantially improved average signal-to-noise ratio (SNR) across the entire q-range, notably in the water scattering region. At q ≈ 3.8 Å^−1^, the average SNR in the aerosol SFX experiment reaches ∼580, whereas that in the liquid-jet based SFX experiment is only ∼1.3. The significantly reduced background in aerosol SFX enhances the average SNR and enables more precise measurements of Bragg diffraction from weakly diffracting nanocrystals.

### Aerosol SFX structure

We chose CpGV granulin nanocrystals as a model system to systematically validate the applicability of aerosol delivery-based SFX for several key reasons: i) CpGV granulin nanocrystals are naturally formed in CpGV occlusion bodies without physical interferences, ii) the crystals preserve high homogeneity in both size (500 × 250 × 250 nm^3^) and a spheroid volume (∼0.016 *µ*m^3^), containing only several thousand unit cells per crystal (Supplementary Fig. 2a,b) and molecular content, iii) the crystals can be produced in a high density slurry (1×10^12^ crystals · ml^−1^) [40], and iv) the CpGV nanocrystals can be delivered in extremely low viscous medium, mitigating clogging issues due to high viscousity media [5]. CpGV structure has been resolved using both conventional SFX (liquid jet-based SFX) [5, 14] and a cryogenic serial electron diffraction [41] to 2.1 Å and 1.6 Å, respectively, providing important structural information to benchmark this novel method. Moreover, Awel et al. have previously demonstrated the feasibility of obtaining strong Bragg diffraction beyond 3.0 Å from CpGV nanocrystals delivered using a convergent-nozzle aerosol injector. However, only a limited number of hits were recorded (∼0.006% hit rate) [36]. Note that a photon energy of 8 keV was used with an average pulse energy of ∼2.94 mJ and a focal spot of 1.3 *µ*m × 1.3 *µ*m (H × V, FWHM) at the interaction point, corresponding to ∼2.29 × 10^12^ photons · *µ*m^−2^.

To ensure a high photon flux on the CpGV nanocrystals, we used an unattenuated 6 keV X-ray beam (*λ* = 2.04 Å) focused to a size of approximately 300 × 300 nm^2^ (FWHM) at the interaction region, with a pulse energy of ∼2.5 mJ. This corresponds to a photon fluence of approximately 2.9 × 10^13^ photons · *µ*m^−2^ at the interaction region. Using this configuration, we successfully executed aerosol SFX experiments at the European XFEL, operating at an intra-train repetition rate of 1.128 MHz with 352 pulses per train and 10 Hz train repetition rate with ∼25 fs pulse length, to acquire diffraction data from aerosolised CpGV nanocrystals (Table 1) [42, 43]. This enabled us to process and index 32 031 crystals, with an average hit rate of 0.029% and an indexing rate of 115% of the frames containing hits (Table 2), diffracting to the edge of AGIPD detector with high-resolution Bragg diffraction at 2.26 Å (Supplementary Fig. 2c). Supplementary Fig. 3a,b compares the pseudo-powder patterns (based on peak counts) for overall indexed and non-indexed hit frames with 96.54% and 3.46%, respectively.

**Table 1:**
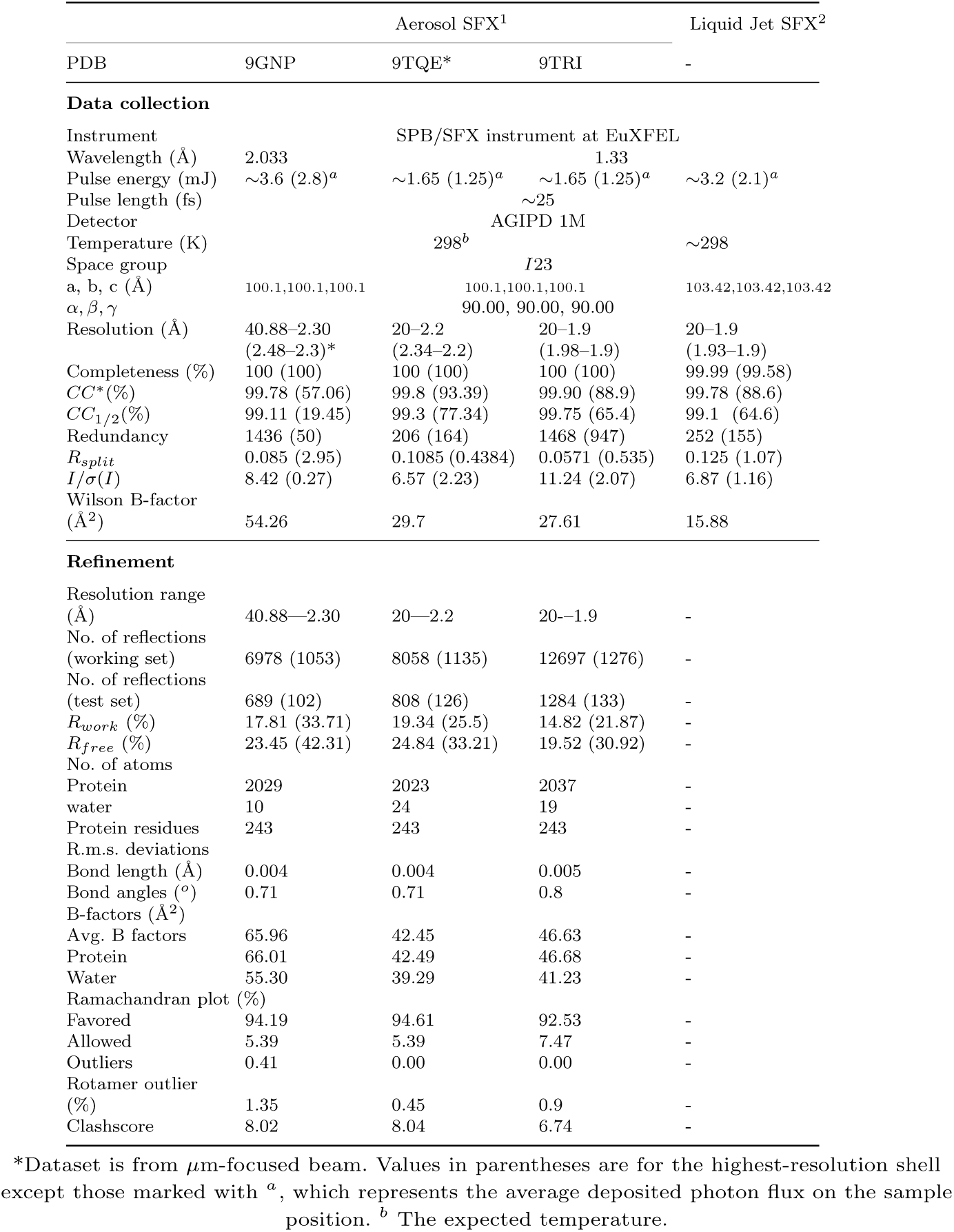
Data collection and refinement statistics.

**Table 2:** Comparison of aerosol and liquid-jet delivery method.

| Experiment | Aerosol 6 keV | Aerosol 9.3 keV<br>(defocused) | Aerosol 9.3 keV<br>(focused) | Liquid jet |
| --- | --- | --- | --- | --- |
| Collection time<br>(min) | 405 | 175 | 524.5 | 12 |
| Total frames | 97 798 624 | 36 838 152 | 109 367 388 | 1 469 109 |
| Crystal hits | 27 961 | 9 082 | 31 204 | 21 784 |
| Indexed<br>crystals | 32 031 | 7 176 | 28 927 | 19 919 |
| Indexing rate<br>(%) | 115 | 79.0 | 92.7 | 91.4 |
| Used CpGV<br>crystals | $5.16 \times 10^{11}$ | $1.61 \times 10^{11}$ | $6.20 \times 10^{11}$ | $1.8 \times 10^{10}$ |

We applied a resolution cut-off of 2.3 Å for intensity merging and resolved, the first aerosol SFX crystal structure (PDB ID: 9GNP) using molecular replacement with a liquid-jet-based SFX structure (PDB ID: 5G0Z or 5MND) as the search model, yielding a model of reasonable quality with *R*_work_ and *R*_free_ of 17.81% and 23.45%, respectively (Table 1). Our diffraction patterns show strong Bragg peaks at the corners and edges of the detector at a photon energy of 6 keV, indicating that higher resolutions can be attainable at higher photon energies and optimised ALS exit orifice alignment. During the 6 keV measurements, a substantial area near the detector edge was shadowed by the shroud, leading to a reduction of ∼0.2 Å in the effective resolution (see Supplementary Fig. 2c).

Comparison of the aerosol SFX model with that of the conventional SFX (liquid jet-based SFX) points to two key related differences: i) shrinking of the unit cells by ∼3.3%, shortening the cubic axes to ∼100 Å (Fig. 3a), and ii) a ∼35% reduction in the solvent content. Such a significant decrease in the solvent content of the unit cell is corroborated by the five-fold reduction of water molecules in the aerosol SFX model. We attribute this to the dehydration effect during aerosolisation, due to evaporation [44, 45], which yielded a 8.33% decrease in the solvent accessible surface area of the aerosol structure in comparison with the liquid jet-based structure, 16 436 Å^2^ and 17 805 Å^2^, respectively. We were able, however, to model some of the water molecules (Table 1). Consistent with previous reports [31, 46, 47], our results indicate that *in vacuo* aerosolisation retains sufficient hydration, thereby preserving the integrity of biological samples, suggesting that the kinetics of the desiccation exceed the transit time of the droplets to the interaction point.

**Fig. 3:**
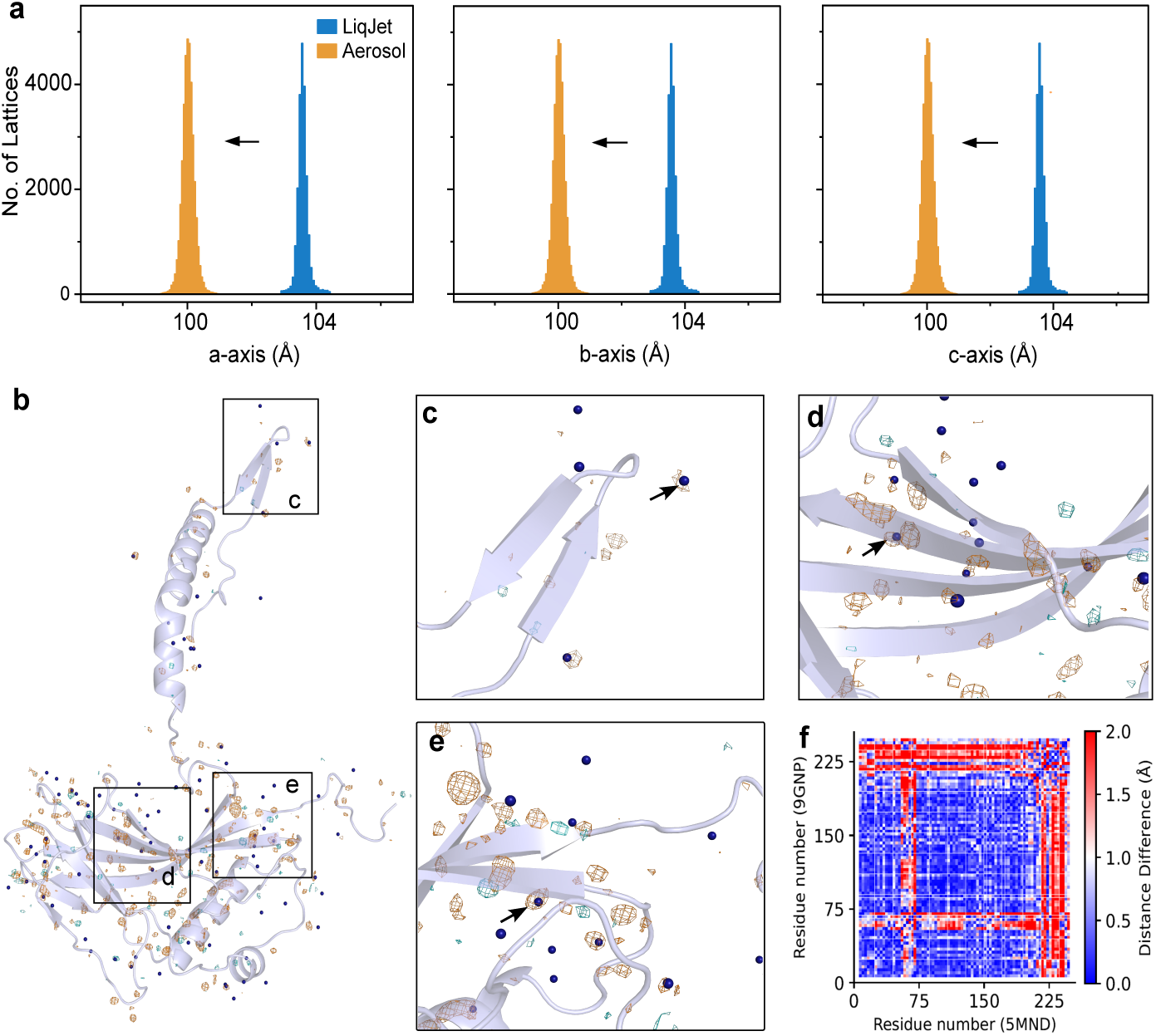
Comparison of aerosol and liquid jet-based SFX. (**a**) Shrinking of the unit cell axes of aerosolised CpGV nanocrystals in comparison with that of the liquid jet (PDB ID: 5MND) as indicated by the arrows, (**b**) *F_O_^aerosol^* − *F_O_^liqJet^* isomorphous difference map displaying the positive (cyan mesh) and negative (orange mesh) peaks contoured at 3.0 sigma level, (**c**) selected site in the N-terminal, (**d**) loop, and (**e**) the C-terminal region, respectively, showing the disappearance of the water molecules in the aerosol SFX structure as indicated by the negative peaks. The arrows in (c,d,e) show some of the negative amplitude difference maps around water molecules indicative of the disappearance of these waters in the aerosol structures, and (**f** ) the distance difference matrix analysis calculated using the liquid jet-based model (PDB ID: 5MND) and the aerosol model of 6 keV (PDB ID: 9GNP).

We note that adding a diffusion dryer downstream of the nebulization chamber does not alter the dehydration behavior of CpGV nanocrystals. The shrinking of the unit cells led, to some extent, to a lower-quality electron density map, particularly in the N-terminal site, the loop region, and specifically the intrinsic loop formed by residues 175–190 at the core of the structure. These domains were found to be challenging for a synchrotron-derived structure, owing to their high mobility [5], suggesting also a potential effect of X-ray radiation damage.

To compare the two structures, we next calculated the 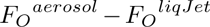 isomorphous difference electron density map, using the liquid jet-based model (PDB ID: 5MND) as a reference (Fig. 3b). Intriguingly, the difference map reveals strong negative peaks (≥3 *σ* level) across the aerosol structure, which was clearly absent in the 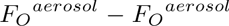 isomorphous difference maps (Supplementary Fig. 4). A large number of these negative peaks were water molecules (Fig. 3c–e), consistent with the observed shrinking trend. This suggests that aerosol sample delivery under vacuum induces dehydration of the unit cells, leading to a reduction in the overall solvent molecules, due to tightening of crystal contacts [48]. This was confirmed by the distance difference matrix analysis between the liquid jet and the aerosol based model, which revealed significant shifts (up to 4 Å) at several regions in the aerosol structure, especially residues 60–75 (the solvent exposed loop) and residues 210–225 (the C-terminal region), which together contribute to the polyhedral assembly (Fig. 3a,f). These domains play central role in conformational plasticity in viral polyhedral structures, consistent with the localised increase of B-factor relative to other regions of the protein [5, 49, 50].

### High-resolution structure of aerosol SFX

To further validate these observations, we then used 9.3 keV X-ray photon energy with both defocused micrometre-scale (2.5 × 2.5 *µm*^2^) and nm-focused X-ray beams to collect high-resolution diffraction data from aerosolised CpGV nanocrystals for low and high photon fluxes, respectively (Table 1). This resulted in Bragg diffraction peaks beyond 2.0 Å and 1.7 Å resolution, respectively (Supplementary Fig. 2d). The pseudo-powder pattern over the total indexed (∼90%) and non-indexed hit frames (∼10%) are similar to that obtained for the 6.0 keV data (Supplementary Fig. 3), suggesting consistency in aerosol SFX experiments at different X-ray energies. For comparison, we collected liquid jet-based SFX data on CpGV nanocrystals using a DFFN nozzle at 9.3 keV, diffracting to 1.9 Å resolution. We then compared the figure of merit (FOM) of these data at a resolution cut-off of 2.3 Å, which revealed significant similarities between different datasets, except that the aerosol SFX data yielded significantly higher SNR in comparison with that of the liquid and aerosol SFX with a defocused beam (Supplementary Fig. 5).

We solved the structures (9.3 keV) at a resolution of 2.2 Å and 1.9 Å, respectively (Table 1). The C-*α* least square superimposition of the two structures resulted in a root mean square deviation (rmsd) of 0.125 Å, indicating high similarity between the two structures, similar to that of the 6 keV model, which gave rise to an rmsd of 0.17 Å when superimposed with the 9.3 keV models. Thus, we focused our analysis on the high-resolution data, which yielded improved statistics at the cut-off bin of 1.9 Å resolution with a correlation coefficient (*CC*_1*/*2_) of 66.4% and refined to an overall *R*_work_ and *R*_free_ of 14.82% and 19.52%, respectively, indicating intrinsic consistency (Supplementary Fig. 6a–e).

Figures 4a–e show the homo-dodecameric biological assembly, the monomeric overall architecture of the 1.9 Å resolution model, and the quality of the electron density maps. Superimposition of the aerosol with the liquid jet structure at 1.9 Å resolution resulted in an rmsd of 0.85 Å, indicating high global similarity, as shown in the 6 keV case (∼1.17 Å to the liquid-jet based model; Fig. 3f), with higher B-factor in the 6 keV model (Supplementary Fig. 7). This indicates that the aerosol crystal exhibits a strong transition into a dehydration state, while preserving the overall structural integrity of the polyhedrin, suggesting systematic rather than random structural changes during aerosolization (Fig. 4f). This is substantiated by the high structural similarity among the aerosol structures with an average rmsd of ∼0.15 Å. The transition from the hydration to the dehydration state likely drives the thermal mobility observed within the core loop and the N-terminal region, as evidenced by the elevation of the B-factor in these specific regions (Fig. 4g,h) [51].

**Fig. 4:**
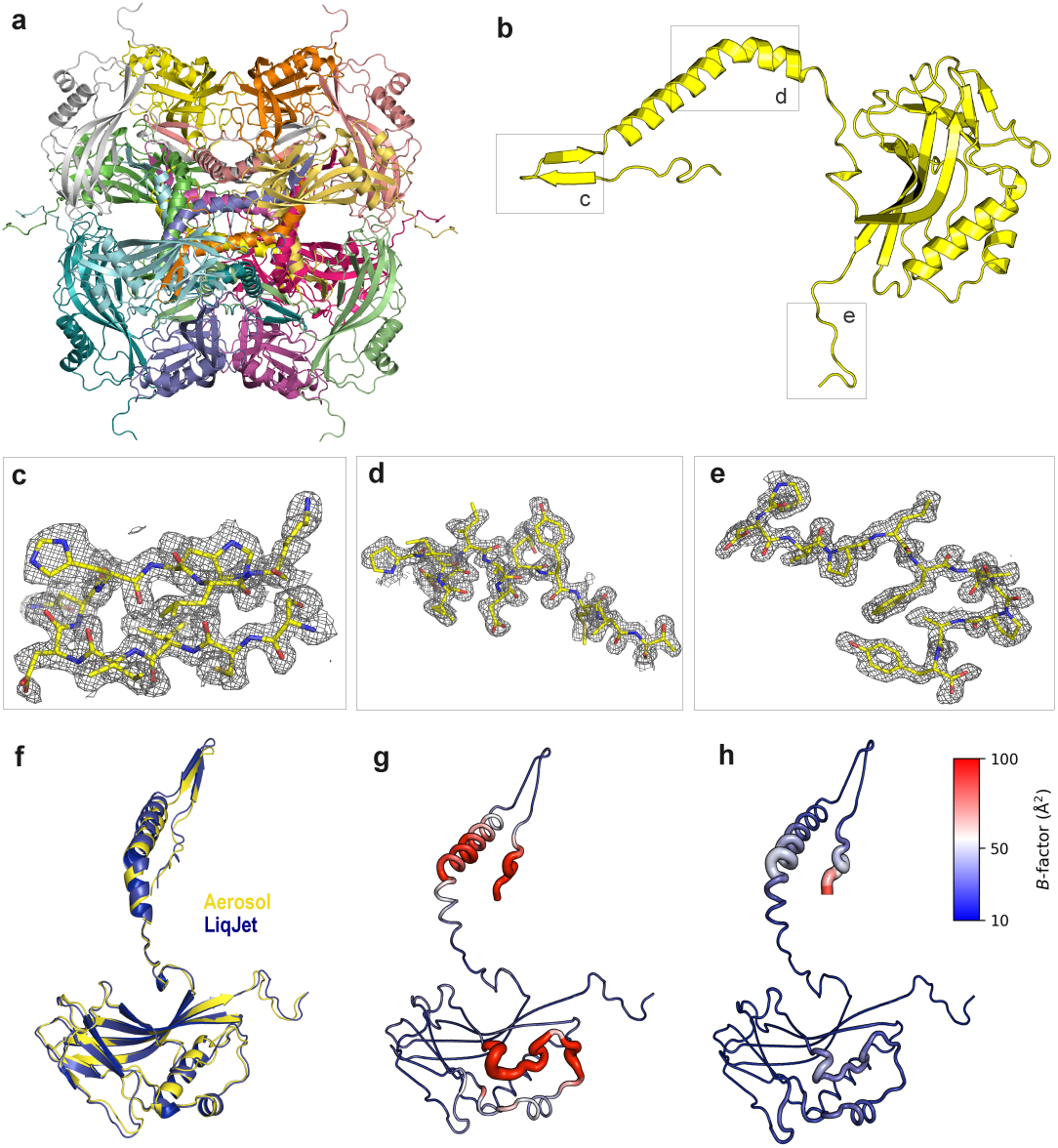
Overall structure of high-resolution aerosol SFX. (**a**) biological assembly (a homo-dodecameric complex) of the high-resolution of CpGV at 1.9 Å, (**b**) monomeric structure of CpGV, (**c**) electron density map quality at the N-terminal region, (**d**) the loop, and (**e**) the C-terminal region, respectively, as indicated in (b). The electron density map (gray mesh) is contoured at 1.0 sigma level. (**f** ) The superimposition of the monomeric structure of the aerosol SFX structure (PDB ID: 9TRI) against that of the liquid jet model (PDB ID: 5MND), (**g**) displaying the cartoon model of the aerosol SFX structure colored by B-factor, and (**h**) is the liquid jet structure coloured by B-factor. The B-factor colouring spectrum range of is composed of blue, white, red (from low to high B-factor within the range of 10–100 Å^2^).

### Radiation-damage in aerosol SFX

To obtain high-resolution structure from nanometer-sized crystals with less than ∼1 *µ*m^3^ volume, a high photon flux is required [5, 52–54]. However, extremely high photon fluxes may introduce local as well as global damage in protein structure even within ∼30 fs pulse duration [27, 55]. It has been shown that a single photoelectron at 6 keV can generate up to 300 secondary electrons through ionization cascades, which is sufficient to cause damage in protein crystals within tens of femtoseconds, particularly in radiation-damage-sensitive regions, such as disulfide bridges and carboxylate [55].

To evaluate this, we calculated the average radiation dose for each CpGV structure using RADDOSE-XFEL [56, 57]. Supplementary Table 2 lists the average doses deposited on the three aerosol SFX structures in comparison with that of the liquid jet SFX. At 6 keV, we estimated the dose absorbed by CpGV nanocrystal to be 18.72 MGy, whereas those at 9.3 keV are 1.152 MGy (nm-focused) and 50.4 kGy (*µ*m-focused), low-dose, in comparison with an average dose of 0.987 MGy (nm-focused) for the liquid jet structure. It was estimated previously that, for room-temperature structure, an average dose limit of 0.4 MGy is important to circumvent local damage in proteins such as the disulfide bond and the carboxylate functional groups [57, 58]. This dose limit can be slightly higher when using the ultrashort pulses (≤25 fs) at XFELs [58]. Therefore, these doses are within the acceptable limit at XFELs with the exception of the 6 keV model that significantly lies above the limit.

The Wilson plot comparison of the intensity between the liquid jet and aerosol data at 1.9 Å resolution clearly shows the rapid steeping of the aerosol data (Supplementary Fig. 6f), likely due to the spike and higher fluctuations in the B-factor of the aerosol data (Supplementary Fig. 7). Together, this may indicate that the high B-factor of this model is likely due to both radiation damage in addition to the aforementioned mobility, owing to the physical force imposed on the crystal unit cell due to rapid dehydration during aerosolisation (Supplementary Fig. 8a–d). These regions with high temperature B-factors, comprising loops (highly mobile), N-terminal helix, and the extension of the C-terminal, serve as active domains for biological assembly and crystal contacts (Fig. 3f and Supplementary Fig. 8e) [49, 50].

### Future direction of aerosol SFX

In the aerosol delivery method, a low salt content in the sample delivery media is preferred, which may hinder to some extent the versatility of the aerosol delivery method as has been shown in aerosol delivery-based SPI experiments [30]. This is especially true for protein crystals that either require high salt content in their crystallization conditions or can be dissolved due to low salt content when transferred into low salts sample delivery. However, it remains a powerful and promising tool for proteins that can be crystallized with the salting-out method, for instance. We also note that dehydration and shrinking of the unit cell up to 10% was a key factor for improving resolution in complex cases such as the cyanobacterial photosystem II, giving rise to a factor of ∼1Å improvement [59, 60]. This suggests that a fine-tuning of the current setup may yield gentle dehydration, thereby improving resolution, while preserving the crystalline intactness and hydration state.

Given further optimisation, the aerosol sample delivery method can also be a method of choice due to its high SNR (Fig. 2c; Supplementary Fig. 5e), owing to its very low to no solvent background scattering (Supplementary Fig. 2c–d), compared to the liquid or the high-viscosity sample delivery media. We note that this current state-of-the-art delivery system is also a potential delivery method for small molecule serial femtosecond crystallography (smSFX), which is essential for large-scale structural determination and characterization of pharmaceuticals and functional materials [61–64]. Thus, it provides a suitable platform for rapid, high-throughput characterization of these materials, given the high repetition rate of the European XFEL facility and high photon energies (∼15 keV) with an average pulse intensity of 500 *µ*J. This is also applicable to future facilities operating at high repetition rates such as LCLS-II and SHINE [65, 66]. Small molecule nanocrystalline suspension with up to ∼ 1 × 10^15^ crystal · ml^−1^ can readily be delivered using our aerosol sample delivery method without further treatment, as most of the small molecule nanocrystallines are compatible with various organic solvents that are suitable for aerosol-based delivery, owing to their volatility [61, 67]. The hit rates can be further improved by controlling the relative concentration of the nanocrystals and by optimising the design of the nebulization chamber and ALS [30, 68]. Unlike nozzle-based liquid jet delivery methods, where the maximum X-ray repetition rate is limited by shockwave propagation in the jet and is directly coupled to the sample delivery rate or consumption rate, the aerosol SFX method is decoupled from sample consumption and therefore represents a promising delivery method for operation at higher repetition rates XFELs (Fig. 2d)[69, 70].

It is noteworthy that the dehydration effect we observed in the protein crystals can significantly improve crystal packing, and is hence expected to further enhance the quality of smSFX data [71]. Notably, water is not an essential factor to maintain the durability of the small molecule materials within the nanocrystalline materials [71, 72]. Therefore, we anticipate that the aerosol sample delivery method can advance the field of smSFX significantly, paving the way also for time-resolved SFX applications in studying, e.g. the non-equilibrium behaviour of nanocrystallines, phase transition and light-induced charge-transfer in crystalline materials with small lattices or novel crystalline materials, such as metal organic frameworks (MOFs) [64, 67, 73–75].

## Discussion

We demonstrate the successful use of the aerosol sample delivery method in protein SFX at a high-repetition-rate XFEL. This approach enabled high-resolution structure determination of CpGV granulin nanocrystals containing only a few thousand unit cells. While both methods (liquid jet and aerosol) yield high-quality structural data, the aerosol model exhibits reduced solvent content, owing to the rapid evaporation during aerosolisation. Importantly, we show that the aerosol delivery method can provide an enhanced SNR by minimizing background scattering similar to previous reports [36], which is crucial for time-resolved experiments, notably pump–probe SFX, as it improves excitation laser penetration depth into crystalline samples [55, 76]. As such, this method provides a unique platform for investigating, e.g. ultrafast photocatalytic mechanisms and crystalline phonon dynamics in small lattices, where high SNR is indispensable for capturing subtle structural fluctuations [77]. We anticipate that these advantages can be further exploited in smSFX for the high-throughput structural characterization of pharmaceuticals and diverse crystalline materials, e.g. MOFs and crystalline inorganic transistors [61, 67]. In summary, this method opens new avenues for SFX studies of functional materials and macromolecules that remain stable in low-salt mother liquors. Beyond crystallography, this method serves as a methodological bridge to SPI. By probing the diffraction limits of nanocrystals containing only a few unit cells, approaching the single-molecule regime, aerosol SFX provides empirical constraints for calculating the dose and particle-size requirements crucial for pushing the SPI towards sub-nanometer resolution [78–81]. Furthermore, the low solvent background in aerosol SFX enables hit finding based on total scattering signals, thus simplifying data reduction workflows [82, 83].

## Methods

### Preparation of CpGV nanocrystals

CpGV was purified from the commercial formulation Madex TOP (BioFA). 10 ml of Madex TOP was diluted with 40 ml of ultrapure water and distributed into two 50 ml centrifuge tubes. Samples were centrifuged for 10 minutes at 10 000 × g at room temperature. The supernatant was discarded, and the pellet was resuspended in a total volume of 25 ml of ultrapure water. The suspension was passed through a 100 *µ*m gravity filter (CellTrics, Sysmex) and allowed to sediment overnight. After removal of the supernatant, the pellet was resuspended in 15 ml of ultrapure water and sequentially filtered by gravity through 100, 50, 30, 20, 10, and 5 *µ*m filters (CellTrics, Sysmex). Particle concentration was assessed by nanoparticle tracking analysis (NanoSight NS300, Malvern Panalytical), yielding a final concentration of 2.5 × 10^11^ CpGV particles · ml^−1^ in a total volume of 10 ml. Purified virus preparations were stored at 4^◦^C until further use.

### Sample delivery

The sample delivery system consists of a GDVN or DFFN for sample aerosolisation, a differential pumping (skimmer) section, and ALS [38]. The suspension of CpGV nanocrystals is loaded into a reservoir to maintain in homogenous suspension using an anti-settler device. The suspension was aerosolised using a GDVN (for the 6 keV experiment, MVED F-type [70]) or a DFFN (for the 9.3 keV experiment, MVED N-type [84]) with liquid flow rates of 2-5 *µ*l · min^−1^ and helium as a sheath gas. For the DFFN, the second liquid was ethanol with a flow rate of 3-4 *µ*l · min^−1^ (see Supplementary Table 1). The generated aerosol, which is composed of helium and the droplets containing CpGV nanocrystals, passes through a conductive tubing into the differential pumping section. The majority of the droplet evaporation happens up to this section. To assess the impact of dehydration on a sample, a diffusion dryer (Model 3062-NC, TSI Inc.) was incorporated into the conductive tube for the 9.3 keV experiments. The differential pumping section consists of a nozzle-skimmer pair at a variable distance with pumping in between. The remaining gas and the CpGV nanocrystals that pass through the skimmer enter the ALS with a pressure between 1.2–1.9 mbar (injector pressure). The nanocrystals exiting the ALS and entering the interaction chamber are monitored via optical scattering [38]. The pressure inside the interaction chamber is kept at 2.3 × 10^−5^ mbar. The 532 nm laser for particle scattering microscopy arrives 1 ms after the X-ray pulse train to avoid interference with the incident X-ray pulse [85]. The scattering of the CpGV particles can be observed during the experiment on a side-view or in-line microscope and functions as a valuable tool during data collection to ensure a working sample delivery system or before the data collection to align the nozzle-skimmer pair for hit rate and pressure level optimisation. An example of the optical scattering signal on the in-line microscope is shown in Supplementary Fig. 1d.

### Estimation of CpGV particle beam width and velocity

The CpGV beam width estimation is based on particle trajectory calculations. The gas flow field of the ALS is calculated using COMSOL Multiphysics [86], and the particle trajectories are calculated using the Python package CMInject [87]. The helium gas flow field is solved for the upper and lower limit of the experimental injector pressure (pressure above the ALS) of 1.2 and 1.9 mbar in the TSI-AFL100 geometry. The region outside of the ALS after the exit aperture is calculated in a circular piece with the radius of the exit aperture with a pressure boundary condition of 10^−2^ mbar. For the flow field, particle trajectory calculations are performed with the following input parameters: particle density *ρ*= 1.16 g · cm^−3^ [88], an aerodynamic diameter of 320 nm, based on the size of CpGV determined by TEM (500 × 250 × 250 nm^3^). The number of particles is 1000 and Brownian motion is included for the calculation. Simulated particle trajectories inside the ALS are shown in Supplementary Fig. 1a for particles at different radial starting positions. Supplementary Fig. 1b shows the axial velocity of the particles inside the ALS, reaching the maximum velocity outside the ALS. The expected particle beam parameters are between the results of the lower and upper limit of the pressures used and are shown in Supplementary Fig. 1c. We expect a focused particle beam with a diameter of less than 10 *µ*m (*d*_90_) at a distance of 4.2 mm from the ALS exit. The average velocity of the particles is 43 m · s^−1^ for 1.2 mbar and 58 m · s^−1^ for 1.9 mbar inlet pressure at the end of the calculated flow field.

### Data collection

The aerosol SFX experiment was performed at the SPB/SFX instrument [42] of the European XFEL [9]. The machine was operated at a 1.128 MHz intra-train repetition rate with 352 pulses per train mode and an inter-train repetition rate of 10 Hz. We conducted two separate experiments with an aerosol delivery system, using either 6 keV or 9.3 keV X-ray pulses, which were focused by a nano-focusing Kirkpatrick-Baez (KB) mirror system [89] to a beam size of approximately 300 *nm* (FWHM) in both the horizontal and vertical directions or a defocused micrometer-scale beam with FWHM of 2.8 *µ*m [42]. The averaged X-ray pulse energy was about 3.6 mJ at 6 keV and 1.65 mJ at 9.3 keV, measured by X-ray gas monitors in the tunnel [90]. For the liquid jet delivery system, the beam was delivered with 0.564 MHz of intra-train repetition rate and 10 Hz of inter-train repetition rate. The photon energy was 9.3 keV. The diffracted X-ray by each nanocrystal was measured with 1 megapixel adaptive gain integrating pixel detector (AGIPD 1M) [91], located about 125 mm downstream from the interaction point and the direct beam passes through the central gap of the AGIPD 1M detector.

### Data processing

The AGIPD geometry was refined and optimised using powder diffraction and preliminary indexed hits from the granulin nanocrystals with the geomoptimiser programme [92] of the crystFEL software (v9.0.1) [93] to find the beam centre and define the approximate camera length. Online data evaluation was performed with Karabo and OnDa monitor to observe the hit rates, an approximate resolution, integrated intensity (detector saturation), and individual diffraction patterns [94]. A diffraction pattern was considered a hit when it contained at least 10 Bragg peaks. For the offline data processing, the programme crystFEL (v0.11.1) was used via our semi-automated EXtra-Xwiz pipeline [95]. The programme peakfinder8 [96] was used for hit finding with the following peak detection parameters: –peaks=peakfinder8 –min-snr=4 –threshold=5 –min-pix-count=2 –max-pix-count=20 –int-radius=3,5,7 –local-bg-radius=5 –tolerance=15,15,15,5 and –min-peaks=10.

The Xgandalf indexing algorithm [97] was used for the indexing of Bragg peaks from the 9.3 keV data and mosflm [98] for the 6 keV data after initially screened several indexing algorithms, including mosflm, Xgandalf, asdf, and DirAx [99]. The CpGV granulin nanocrystals belong to the spacegroup *I* 23 with two different possibilities in the unit cell axes, which leads to an indexing ambiguity. To resolve this ambiguity, the output stream file of the indexed hits was used as an input for the ambigator tool in the crystFEL software, which was executed with the following re-indexing parameters: -y m-3 -w m-3m and –iterations=4, where the m-3 and m-3m point groups represent the actual symmetry (-y) and the apparent symmetry (-w), respectively [100]. The output stream file of ambigator was then used for scaling and merging partial reflections with the programme partialator of crystFEL (v0.11.1) using the following options: –model=unity -y m-3 –polarisation=horiz –min-measurements=2 – max-adu=inf –push-res=inf and –iterations=3. In total, 32,031 patterns were indexed for the 6.0 keV diffraction data and 36,103 patterns for the 9.3 keV data.

We note that the data did not exhibit any sign of saturation with maximum analogue-to-digital units (ADU) lying well below the saturation level of the AGIPD 1M detector, as has been confirmed by the peakogram calculation via OnDa; therefore, we merged all reflections with the option –max-adu=inf.

Finally, the merged intensity was converted to a structure factor with get hkl, and all FOMs were generated with check hkl and compare hkl of the crystFEL software. Resolution cut-off was applied when the Pearson correlation coefficient (*CC*^∗^) fell below 50%. CpGV nanocrystals diffract to the edges and corners of AGIPD detector at both photon energies; however, to maintain reasonable quality, a cut-off at 2.3 Å and 1.9 Å was applied for 6 keV and 9.3 keV data, respectively.

### Structure determination and refinement

Initial models were obtained by molecular replacement using the molrep programme [101] of the CCP4 suite [102] with the SFX granulin structure (PDB ID: 5G0Z or 5MND), obtained from liquid jet serial crystallography, as the search model. The resulting model was improved manually using the programme Coot (v0.9.8.93) [103]. Rigid body refinement was applied to the model for several cycles using refmac5 with the riding hydrogen model [104]. This was followed by atomic coordinates and temperature factor refinement for several cycles with refmac5. The final rounds of refinement were done using phenix.refine [105] iteratively with manual correction in Coot. Finally, several cycles of Translation-Libration-Screw-rotation (TLS) refinement in PHENIX were applied. The final model geometry was validated with the programme MolProbity [106], and the biological assembly was generated with PISA [107] within Coot. All figures were generated using Pymol [108].

To calculate the 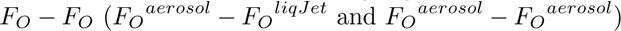 isomorphous difference maps, structure factor amplitudes of the two respective data were scaled using SCALEIT [109] of the CCP4 package to alleviate variation between the two structure factors before calculating the difference map with a peak detection threshold set to a 3.5*σ* level to avoid noise inclusion to the map. Due to its high isomorphism to the aerosol models (rmsd=0.85 Å), the liquid jet model (PDB ID: 5MND) was used as a reference model. For the aerosol isomorphous difference map an aerosol model was used as a reference.

### Calculation of radiation dose

We used the XFEL–customized version of the programme RADDOSE-3D (RADDOSE-XFEL) [56] to estimate the radiation dose absorbed by CpGV nanocrystals. An average nanocrystal spheroid volume of 500 × 250 × 250 nm^3^ and a cubic unit cell axes (a=b=c) of 100.1 Å and 103.41 Å for the aerosol and the liquid jet model, respectively, were used for the calculation with 15 pixels · *µ*m^−1^ and 500,000 photons. The angular resolution was not considered for the calculation since the crystals in serial crystallography are not rotated (wedge angle is 0^◦^). Each calculation was repeated at least three times in triplicates.

## Data availability

Atomic coordinates and structure factors are deposited to the wwPDB and publicly available as: 9GNP: the CpGV model from 6 keV data (https://doi.org/10.2210/pdb9GNP/pdb), 9TQE: the CpGV model from 9.3 keV *µ*m-focused beam (https://doi.org/10.2210/pdb9TQE/pdb), and 9TRI: the CpGV model from the 9.3 keV nm-focused beam (https://doi.org/10.2210/pdb9TRI/pdb). The SFX diffraction data recorded for the 6 keV and 9.3 keV experiments at the SPB/SFX instrument of the European XFEL are available at (https://dx.doi.org/10.22003/XFEL.EU-DATA-900260-00) and (https://dx.doi.org/10.22003/XFEL.EU-DATA-009036-00), respectively.

## Code availability

CrystFEL is available at https://gitlab.desy.de/thomas.white/crystfel under the terms of the GNU general public license. The code used for raddose-xfel calculation is openly available at GitHub: https://github.com/GarmanGroup/RADDOSE-3D.

## Acknowledgments

We thank Dominik Oberthuer (CFEL) for his assistance with the preparation of Granulovirus and Tomas Popelar (European XFEL) with the preparation of the laser system for particle illumination. We acknowledge the sample environment and characterization (SEC) staff of the European XFEL for their assistance with the sample preparation and sample delivery tests prior to beamtime. We are grateful to the SPB/SFX instrument and Data analysis staff for the technical and analytical support. We acknowledge Grant Mills, Seungyeon Hong, and Sunghun Lee for supporting the beamline operation. We acknowledge European XFEL in Schenefeld, Germany, for provision of X-ray free-electron laser beamtimes at SPB/SFX SASE1 under proposal numbers 9036 and 900260, and would like to thank the staff for their assistance. The liquid jet data used here was retrieved with permission from Dominik Oberthuer (main proposer) during beamtime under proposal number 2819. We thank Kurt Ament (European XFEL) for critical proof-reading of the manuscript.

## Author contributions

LW, RS, SK, and JE contributed equally to this work. YK, RdW, JB, and APM initiated the project. YK, FK, RdW, and CK conceptualized and led the project. JS, APM, and RB provided support for the beamtime. RS, RdW, and HH prepared CpGV nanocrystals and other test samples, and RS performed electron microscope characterization of the CpGV samples. YK, LW, and JB prepared and optimised the sample delivery setup with support from RdW, AW, MK, KD, and HH. LW performed COMSOL simulations to estimate the size and velocity of the CpGV particle beam. JK, RL, and TS prepared the nanosecond laser system for particle-scattering microscopy and operated the laser system. YK, FK, RdW, LW, DM, CW, QX, SH, SL, AR, MS, RL, RB, JB, and CK performed the aerosol SFX experiment, including data collection. SK, RdW, FK, ES, and OT performed preliminary SFX data analysis, including online evaluation, geometry preparation, and hit finding. FK performed all formal analysis of SFX data with support from YK, RdW, SK, JE, ES, OT, and CK. YK, FK, RdW, LW, RS, and CK wrote the original draft. YK, FK, RdW, LW, RS, JK, JE, SK, HH, TS, RB, JB, APM, and CK reviewed and edited the manuscript with contributions from all authors.

## Additional information

Supplementary information is available for this paper.

---

**Supplementary Figure 1:**
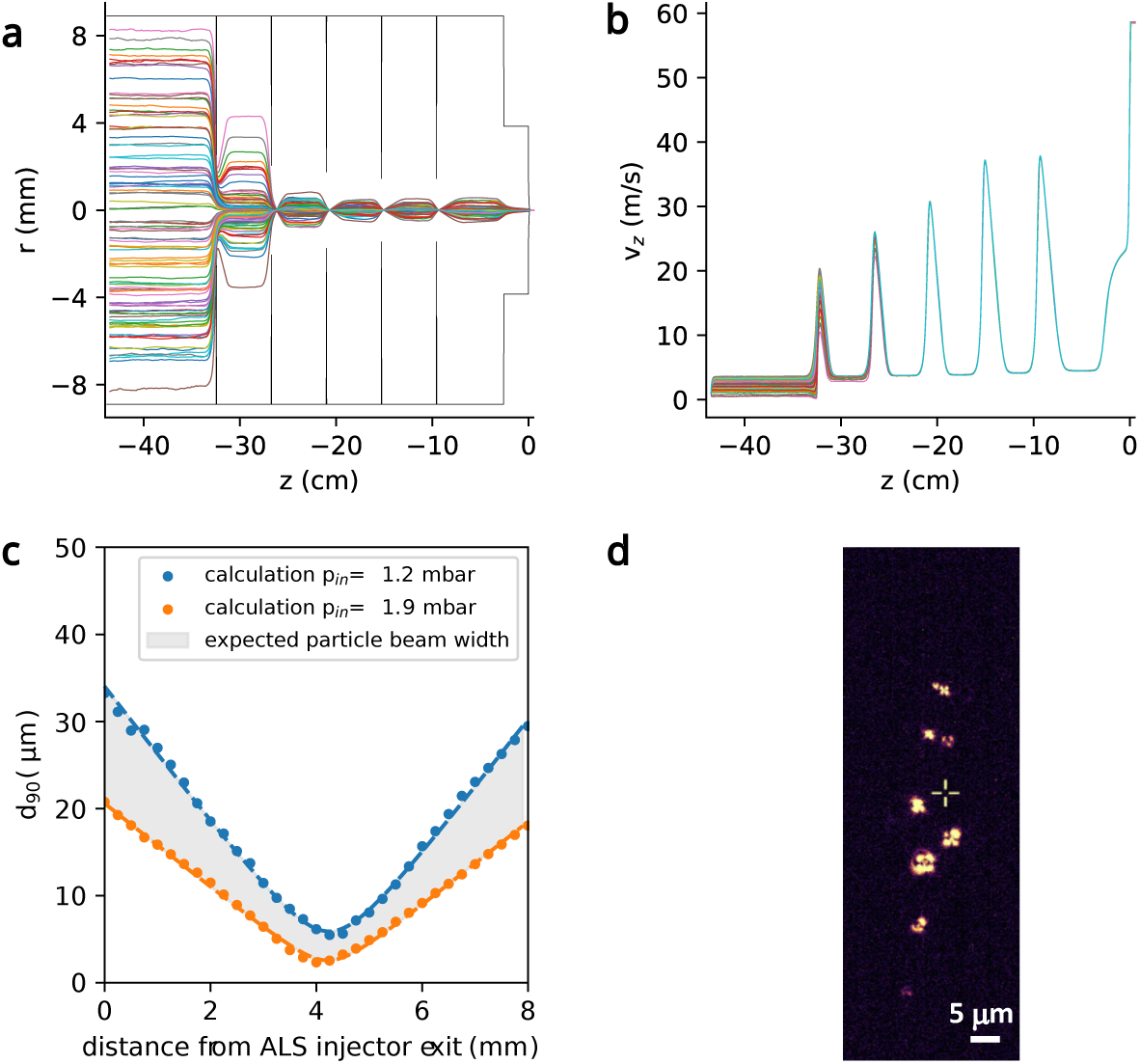
Sample delivery details. Simulated CpGV sample trajectories **(a)** and the particles’ velocities **(b)** from the entrance to the exit of the ALS under 1.9 mbar pressure. The particles were focused and accelerated by passing through the aerodynamic lenses. The speed at the exit of ALS is 58 m · s^−1^. **(c)** Calculated particle beam evolution of the CpGV nanocrystals after leaving the ALS. The gray area represents the expected particle beam size (*d*_90_) at a given distance from the ALS exit. During the experiment, the pressure above the ALS was kept between 1.2 mbar (blue) and 1.9 mbar (orange). The calculated particle beam focus width is below 10 *µ*m at a distance of 4.2 mm. **(d)** A particle scattering image collected during the experiment. Bright spots are the scattering signal of individual CpGV particles in the interaction region illuminated by the 532 nm ns laser. The yellow cross indicates the X-ray beam position.

**Supplementary Figure 2:**
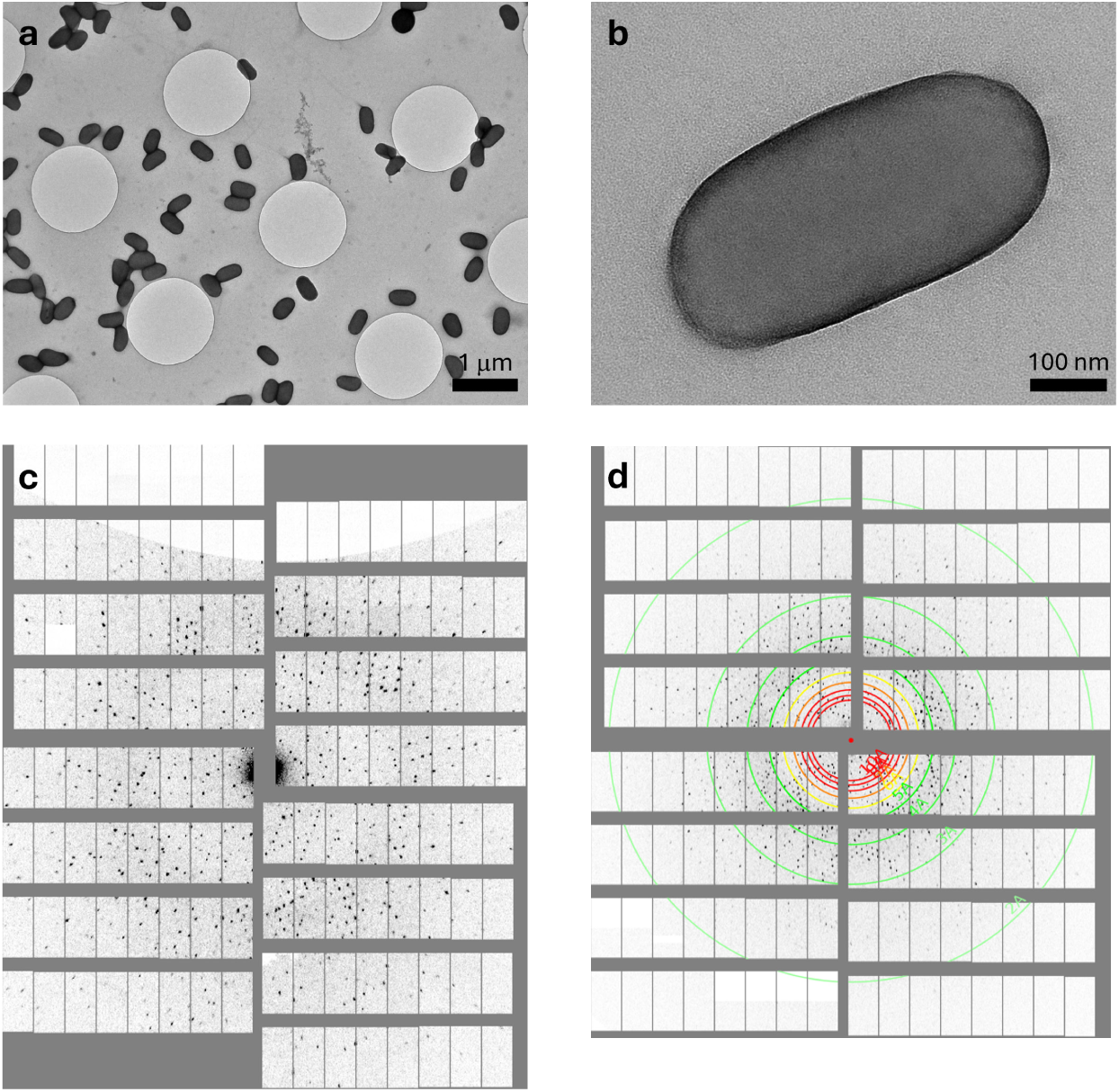
Shape of granulovirus and reperesentative diffraction pattern. TEM image of granulovirus after negative staining with (**a**) low magnification and (**b**) high magnification. Selected diffraction pattern from (**c**) the 6 keV dataset and (**d**) the 9.3 keV dataset.

**Supplementary Figure 3:**
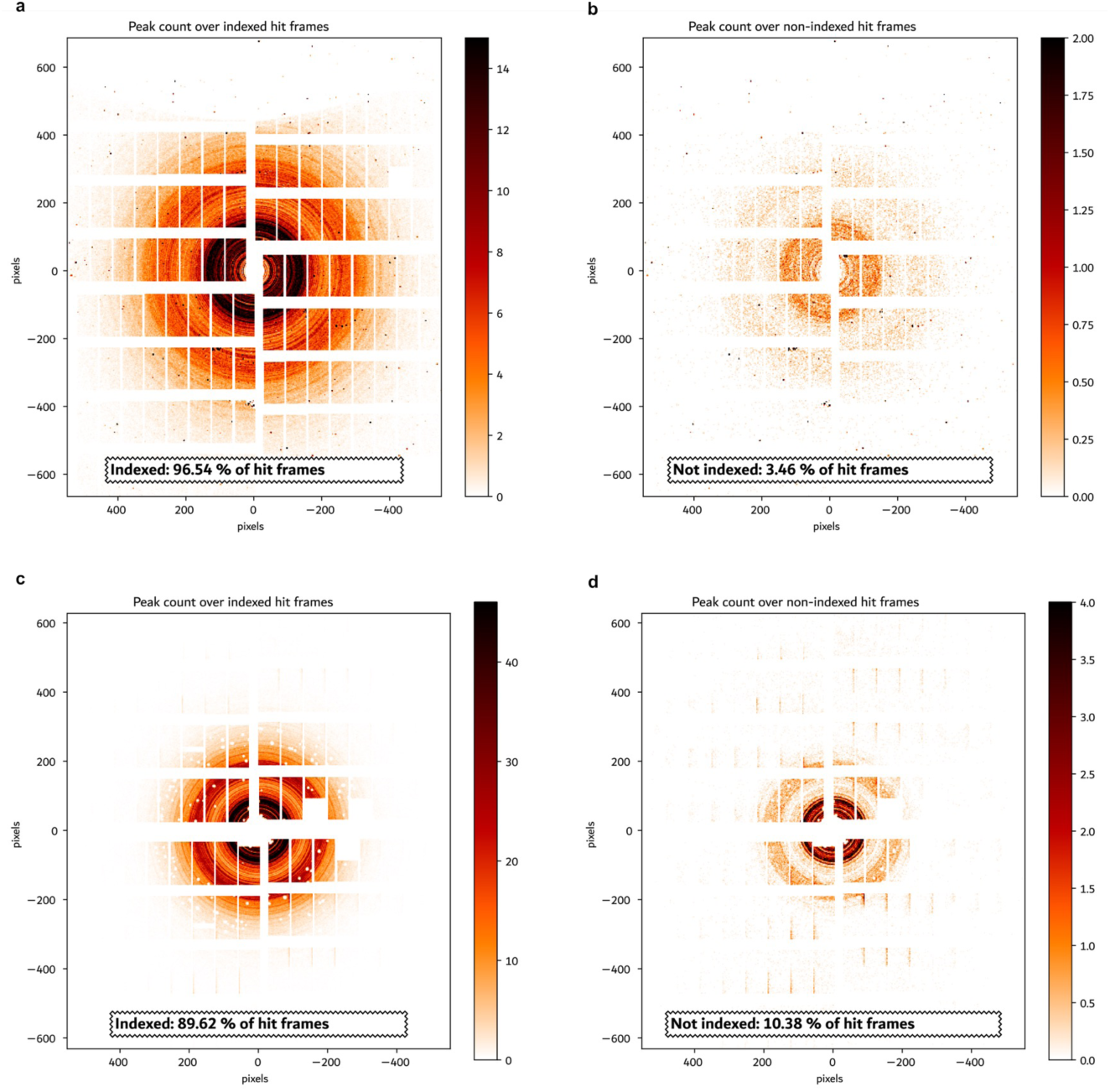
Pseudo-powder pattern from the 6 and 9.3 keV datasets. (**a**) Peak count over indexed hit frames from the 6 keV dataset, (**b**) peak count over non-indexed hit frames from the 6 keV dataset, (**c**) peak count over indexed hit frames from the 9.3 keV dataset, and (**d**) peak count over non-indexed hit frames from the 9.3 keV dataset.

**Supplementary Figure 4:**
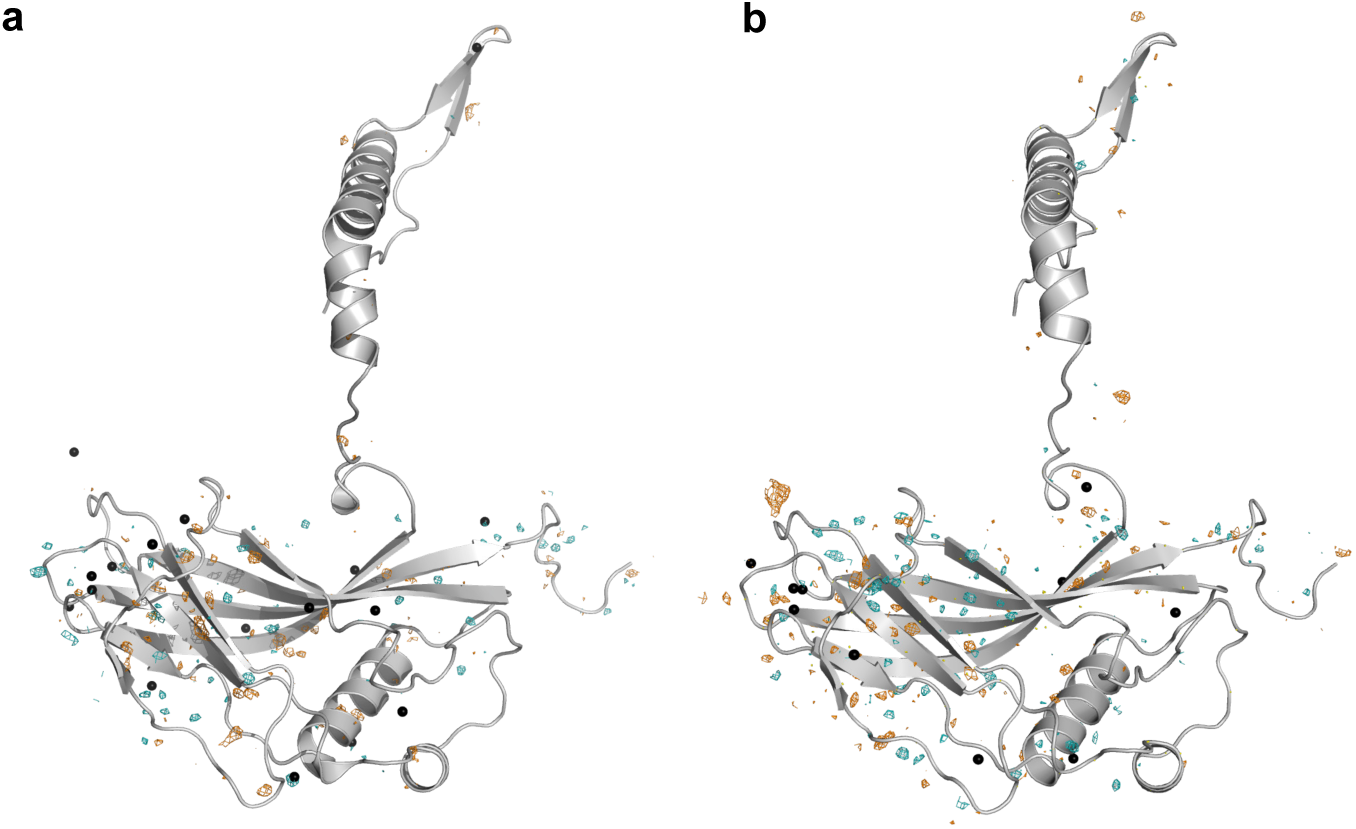
Isomorphous difference maps of the Aerosol structures. (**a**) The isomorphous difference map of 9.3 keV minus that of 6.0 keV scaled at 2.3 Å resolution and (**b**) the difference map of 9.3 keV from the defocused and focused X-ray beams.

**Supplementary Figure 5:**
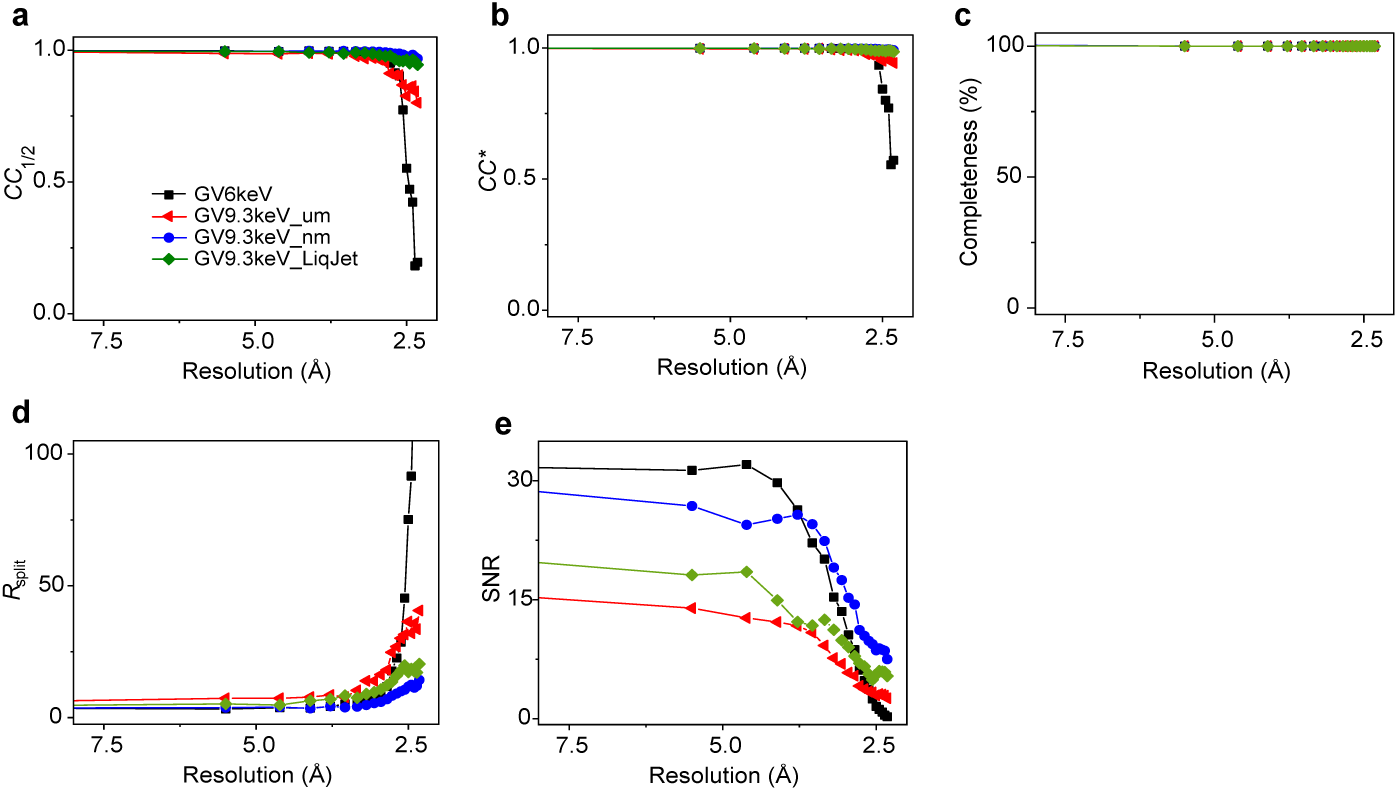
Comparison of the figure of merit of the aerosol and liquid jet with a resolution cut-off at 2.3 Å. (**a**) The *CC*_1*/*2_ Pearson correlation coefficient, (**b**) the *CC*\* correlation coefficient, (**c**) the data completeness, (**d**) the *R*_split_ between odd and even datasets, and (**e**)the signal-to-noise ratios (SNR).

**Supplementary Figure 6:**
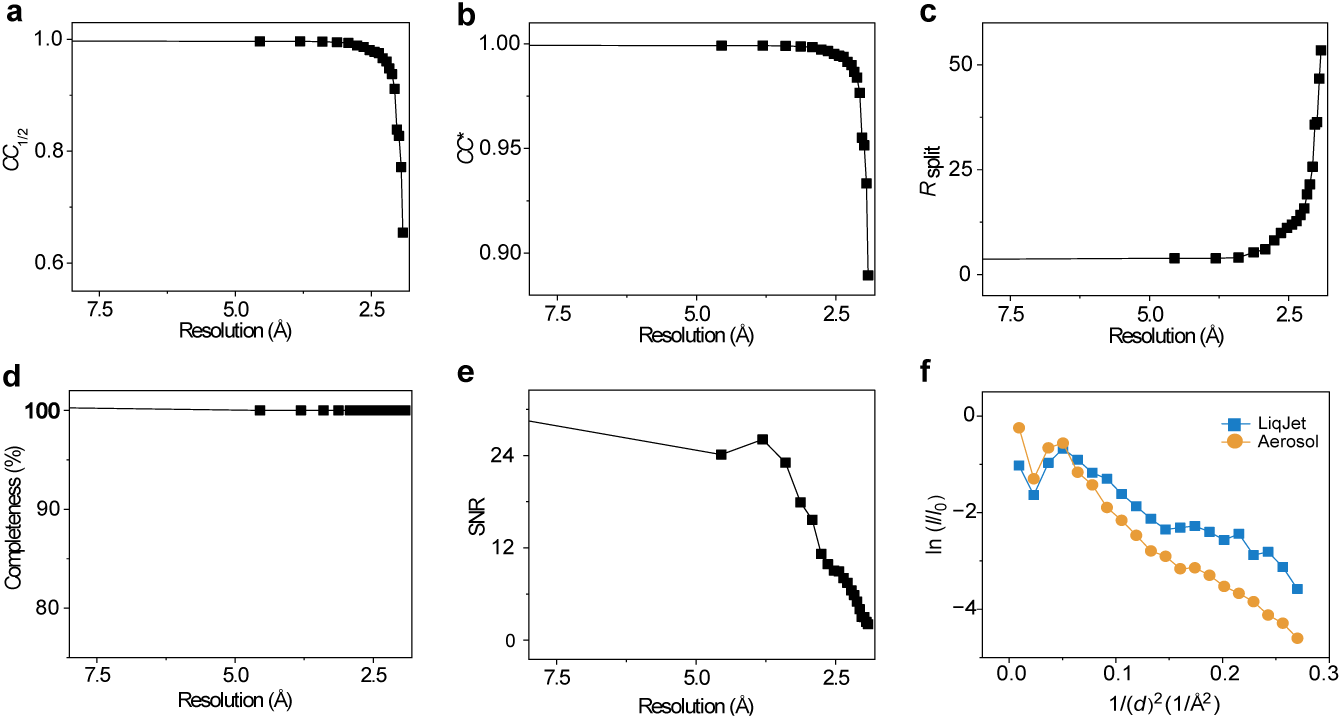
Figure of merit and Wilson plot of the high-resolution aerosol SFX structure resolved at 1.9 Å. (**a**) The *CC*_1*/*2_ Pearson correlation coefficient, (**b**) the *CC*\* correlation coefficient, (**c**) the *R*_split_ between odd and even datasets, (**d**) the data completeness, (**e**) the signal-to-noise ratios (SNR), and (**f** ) a Wilson plot showing the intensity distribution of the high-resolution aerosol dataset compared with that of the liquid jet datasets at 1.9 Å.

**Supplementary Figure 7:**
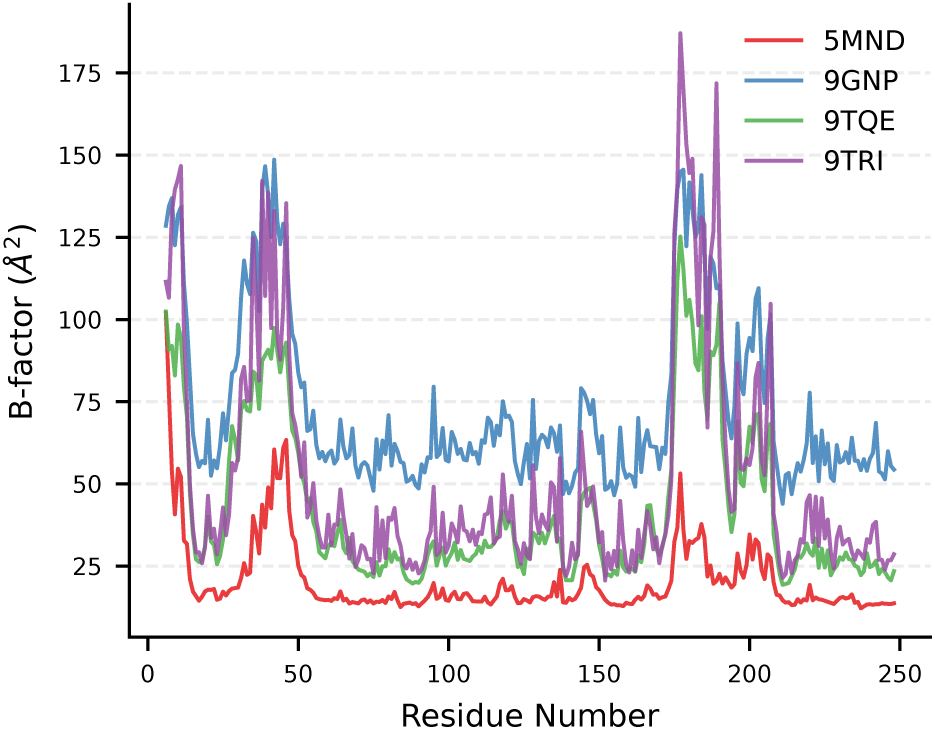
B-factor fluctuations in CpGV structures. The figure shows a comparison of the fluctuations in B-factors across the CpGV structures of aerosl and liquid jet SFX. Three regions—i) residues 6–12 (N-terminal domain), ii) residues 29–50, and iii) and residues 170–210—reveal strong spike in the B-factors of all structures with apparent increase in the aerosol SF.

**Supplementary Figure 8:**
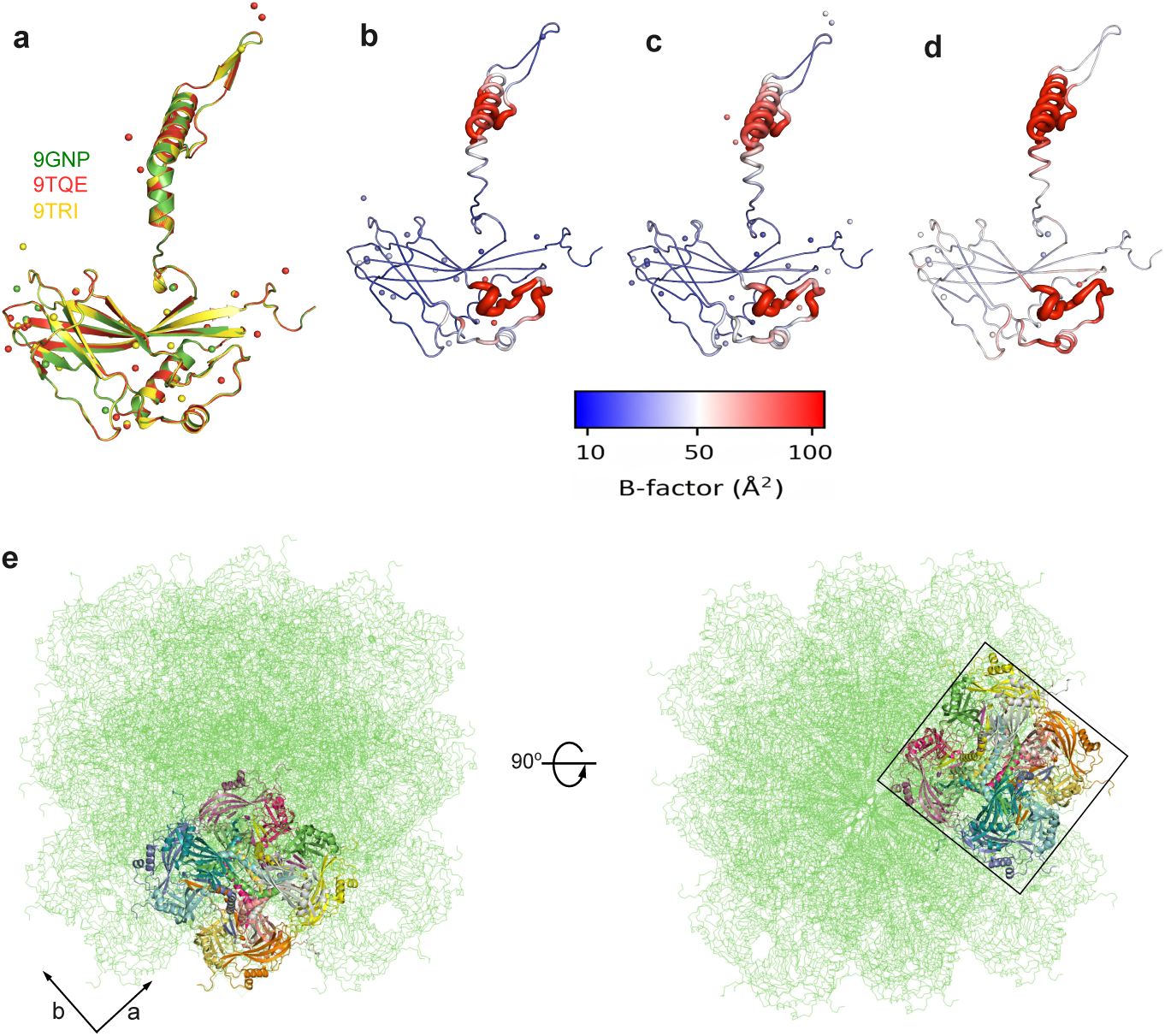
Comparison of aerosol SFX structures. (**a**) The least-square superimposition of the different monomeric structures of the CpGV from different photon and pulse energies (PDB IDs: 9GNP in green (6 keV model), 9TQE in red (9.3 keV with *µ*m-focused beam), and 9TRI in yellow (9.3 keV with nm-focused beam), (**b**) cartoon representation of the 9TQE model coloured based on B-factor fluctuation, (**c**) the cartoon representation of the 9TRI model colored as in (b), (**d**) the 9GNP model colored as in (b,c), and (**e**) crystal packing of the CpGV structure showing 24 symmetry mates at two different orientation with axis a and b shown with arrows and the unit cell phase with squares. The bar in (b–d) represents the B-factor colouring spectrum scale of blue, white, red (from low to high B-factor within the range of 10–100 Å^2^).

**Supplementary Table 1:** Sample delivery parameters for aerosol delivery and liquid jet delivery.

| Experiment | Sample flow rate ($\mu\text{l}/\text{min}$) | EtOH flow rate ($\mu\text{l}/\text{min}$) | He pressure (psi) |
| --- | --- | --- | --- |
| Aerosol 6 keV | 2, 5 | Not used | 310, 350 (410) |
| Aerosol 9 keV | 3 (15) | 3, 4 (15) | 350 |
| Liquid jet | 15 | 20 | 230 |

Note: The numbers inside brackets are the maximum values used when optimizing the condition.

**Supplementary Table 2:** Average radiation doses deposited on CpGV nanocrystals.

|  | PDB | RADDOSE-EXFEL<br>average dose-exposed<br>region (MGy) | RADDOSE-3D style<br>average dose-exposed<br>region (MGy) | Average dose<br>per crystal*<br>(GGy) |
| --- | --- | --- | --- | --- |
| 6 keV<br>(Aerosol) | 9GNP | 18.722 ± 0.35 | 45.231 ± 0.658 | 187.22 |
| 9.3 keV<br>(Aerosol)** | 9QTE | 0.0504 ± 0.012 | 0.208 ± 0.023 | 0.504 |
| 9.3 keV<br>(Aerosol) | 9TRI | 1.152 ± 0.008 | 5.494 ± 0.108 | 11.5 |
| 9.3 keV<br>(Liquid jet) | n.a.*** | 0.987 ± 0.184 | 2.08 ± 0.396 | 9.87 |

\*An average of total dose absorbed by the corresponding number of indexed nanocrystals used to solve the structure; for the 5MND, we considered 10 000 indexed crystals. \*\*A beam was defocused to $\mu\text{m}$ size. \*\*\*Not applicable.

